# Ticks and tickborne diseases in the upper Midwestern United States: role for citizen science in assessing exposure risk

**DOI:** 10.64898/2026.05.14.724901

**Authors:** Alexandra M. Linz, Carrie Marcis, Charles Payant, Linda Donnerbauer, Amber Donnerbauer, Emily Gruenling, Krystal Boese, Garrett Heuer, Alexis Boehm, Johnny A. Uelmen, Thomas R. Fritsche, Jennifer K. Meece

## Abstract

Tickborne diseases are a significant burden in many parts of the world. In the upper Midwestern United States, Lyme disease is the most common tickborne disease. It is carried by *Ixodes scapularis.* This vector can also transmit the pathogens causing anaplasmosis, babesiosis, ehrlichiosis, and several more tickborne diseases in this region. There is also concern for other tick species, such as *Amblyomma americanum,* that are expanding their ranges northward. We launched a citizen science passive tick surveillance program in 2024 to investigate tick species ranges in the upper Midwest, as well as the pathogens carried by *I. scapularis.* We received over 12,000 ticks in the first two years of this program, primarily from Wisconsin. While we received submissions of adult *A. americanum* outside of their endemic range, we did not see evidence of establishment in our study area. We measured pathogen prevalence in adult female *I. scapularis* (n=707) and observed 51% positivity for *Borrelia burgdorferi,* 9% for *Babesia microti,* 9% for *Anaplasma phagocytophilum,* and 3% for *Ehrlichia muris eauclairensis.* Multiple pathogens were identified in 14% of tested specimens, and significant associations were observed between *B. burgdorferi* and *B. microti,* and *B. burgdorferi* and *E. muris eauclairensis.* Pathogen prevalences varied across time and geography. Our results can begin to inform risk assessment for tickborne diseases in our region.

A non-technical version of this document with interactive maps is available here: https://storymaps.arcgis.com/stories/8008c9d710b5400599f3c6cf88b2c546 Our online data dashboard is available <u>here</u>: <u>redcap.link/TICS</u>

## Introduction

Ticks are the principal carriers of vector-borne diseases in North America. Lyme disease, caused primarily by *Borrelia burgdorferi* on this continent, is the most common vector-borne illness in the United States, and cases continue to rise annually (Mead et al., 2024; Rosenberg et al., 2018). *B. burgdorferi* in North America is carried by *Ixodes scapularis* (deer tick or blacklegged tick) and *Ixodes pacificus* (western black-legged tick). Both species can also carry the causative agents of ehrlichiosis, anaplasmosis, and babesiosis. *I. scapularis* can also transmit Powassan virus, *B. miyamotoi,* and *B. mayonii* in the upper Midwest of the United States. One species, *Amblyomma americanum* (lone star tick), may trigger the allergic reaction to red meat, alpha-gal syndrome (AGS), and is also associated with southern tick-associated rash illness (STARI), a disease of unknown etiology (Riva et al., 2023). Some members of the genus *Dermacentor* can transmit Rocky Mountain Spotted Fever in the southeastern and south central United States, and to a lesser extent in the Rocky Mountain region. The ranges of tick species and the pathogens they transmit are expanding as climate conditions change (Sonenshine, 2018). Meanwhile, land use changes mean that more people in suburban and urban environments are coming into contact with ticks (Diuk-Wasser et al., 2021; Gould et al., 2025). Lyme disease is estimated to have been diagnosed and treated in ∼476,000 patients from 2010-2018, while other tickborne diseases, including babesiois and Powassan virus disease, can be fatal (Corrin et al., 2018; Eilbert & Matella, 2024; Kugeler et al., 2021). Together, these factors make tickborne disease a significant public health concern.

A clear delineation of tick species’ endemic ranges is critical. An understanding of where and how these ranges are changing, including each species’ local pathogen prevalences, will be crucial to an effective public health response to tickborne diseases. However, neither of these pieces of data are straightforward to obtain. Tick collection is most often accomplished by dragging a cloth through tick habitat using standard distances and size of cloth. This method of active surveillance effectively estimates tick population sizes at a specific location, but it is labor intensive and time-consuming, limiting our understanding of geographic scope. Pathogen testing in ticks is performed via quantitative real-time PCR (qPCR), a highly sensitive molecular biology technique that requires a sufficient sample size of ticks to estimate pathogen prevalence in a given geographic area – a number large enough that dragging to obtain these ticks can become prohibitively time-consuming. Combined, these challenges limited more accurate pathogen prevalence estimates from being determined in ticks across broad geographic regions.

One way to effectively collect tick species over a large area is through a citizen science approach. This is a passive surveillance method, meaning that it relies on someone encountering, noticing, and submitting/documenting a tick. It inaccurately reflects the populations of tick species in the environment but is highly effective for surveying large numbers of ticks over wide geographic areas (Eisen & Eisen, 2021). Tick citizen science programs have provided important insights into the distributions of ticks and their pathogens. One such effort in Ontario, Canada, served as an early warning system for the arrival of *I. scapularis* and Lyme disease (Ripoche et al., 2018). Another citizen science effort spanning the United States and sponsored by the Bay Area Lyme Foundation, described the range of *Ixodes pacificus* in the western United States and identified *Borrelia* in ticks from counties where it was not previously thought to be endemic (Porter, Barrand, et al., 2021; Porter, Wachara, et al., 2021). In New York, an interactive dashboard was developed to return county-level pathogen prevalence results to citizen science participants, enabling them to assess their own risk of tickborne disease exposure (Hart et al., 2022).

The upper Midwestern region of the United States is highly endemic for several pathogens carried by *I. scapularis,* particularly in Minnesota, Wisconsin, and Michigan. In this region, Lyme disease, caused primarily by *B. burgdorferi,* is the most common tickborne disease, but anaplasmosis and babesiosis also occur. Ehrlichiosis is found in the upper Midwest and is caused locally by *Ehrlichia muris eauclairensis* in *Ixodes scapularis* rather than *E. chafeensis,* the primary pathogen of ehrlichiosis found in *A. americanum* in the southern United States (Pritt et al., 2017). *B. mayonii,* another causative agent of Lyme, is found rarely in this region, as is Powassan virus (Corrin et al., 2018; Rodino & Pritt, 2022). The other tick species of significant abundance in the upper Midwest is *Dermacentor variabilis* (dog or wood tick), which infrequently carries pathogenic *Rickettsia spp.* unlike in other parts of North America (Hecht et al., 2019).

We launched our citizen science surveillance program, the Tick Inventory via Citizen Science (TICS), in the spring of 2024. Here, we present the results of species distributions for the years 2024 and 2025 of the TICS study along with the pathogen testing results from adult female *I. scapularis* submitted in 2024 (Figure 1). The large sample size provided through the participation of citizen scientists allowed us to detect rare species of ticks not yet established in the upper Midwest. It also meant that we could estimate prevalences of the four most common tickborne pathogens in many parts of Wisconsin; over time, this information will assist individuals, clinicians, and public health departments in assessing tickborne disease risk.

**Figure 1.**
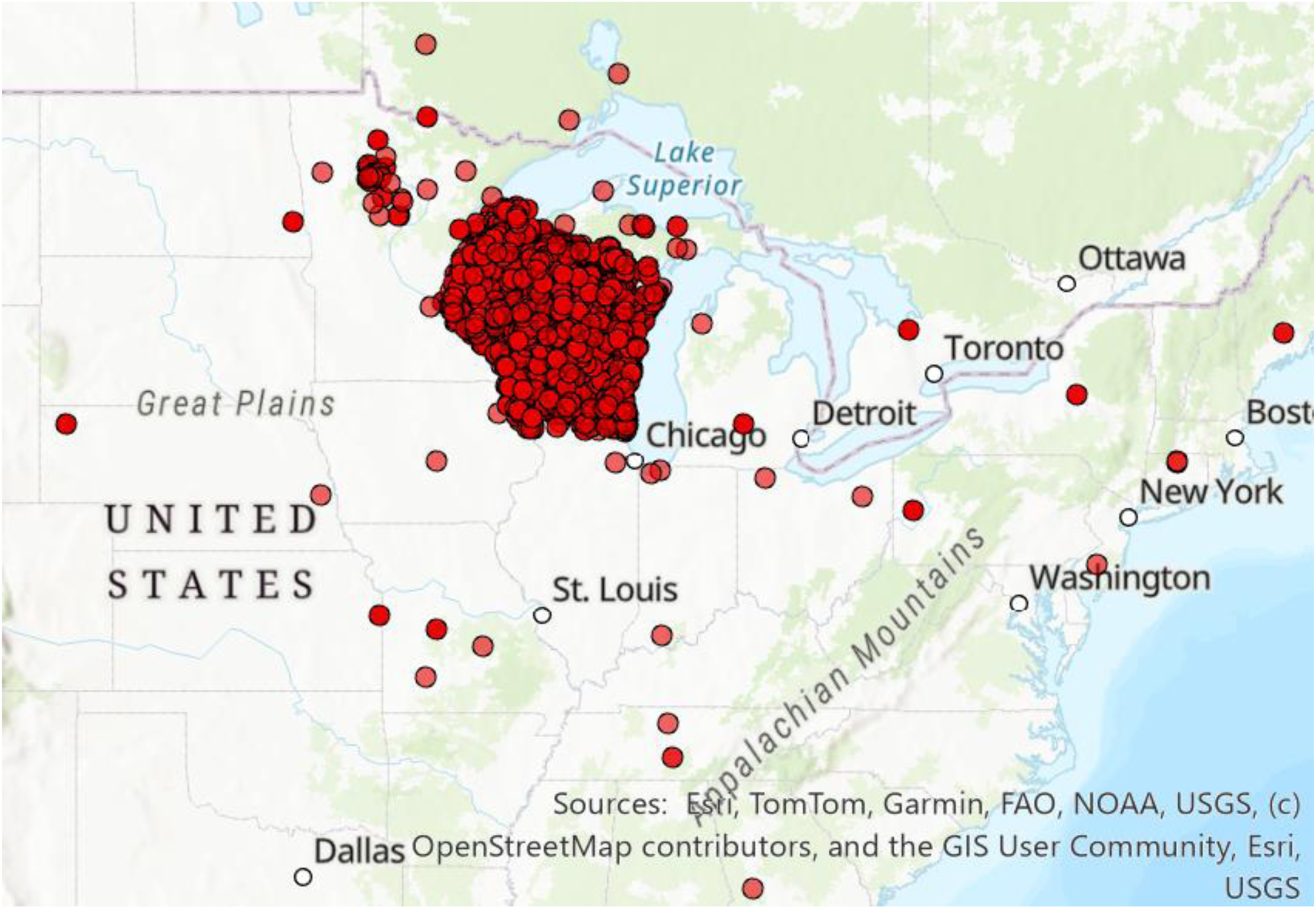
Locations of all submitted ticks in 2024-2025. Kit distribution and advertising was primarily focused on Wisconsin, but we received submissions from across the eastern United States and southern Canada. Each point is semi-transparent, so darker colors represent multiple submissions from the same location.

**Figure 2.**
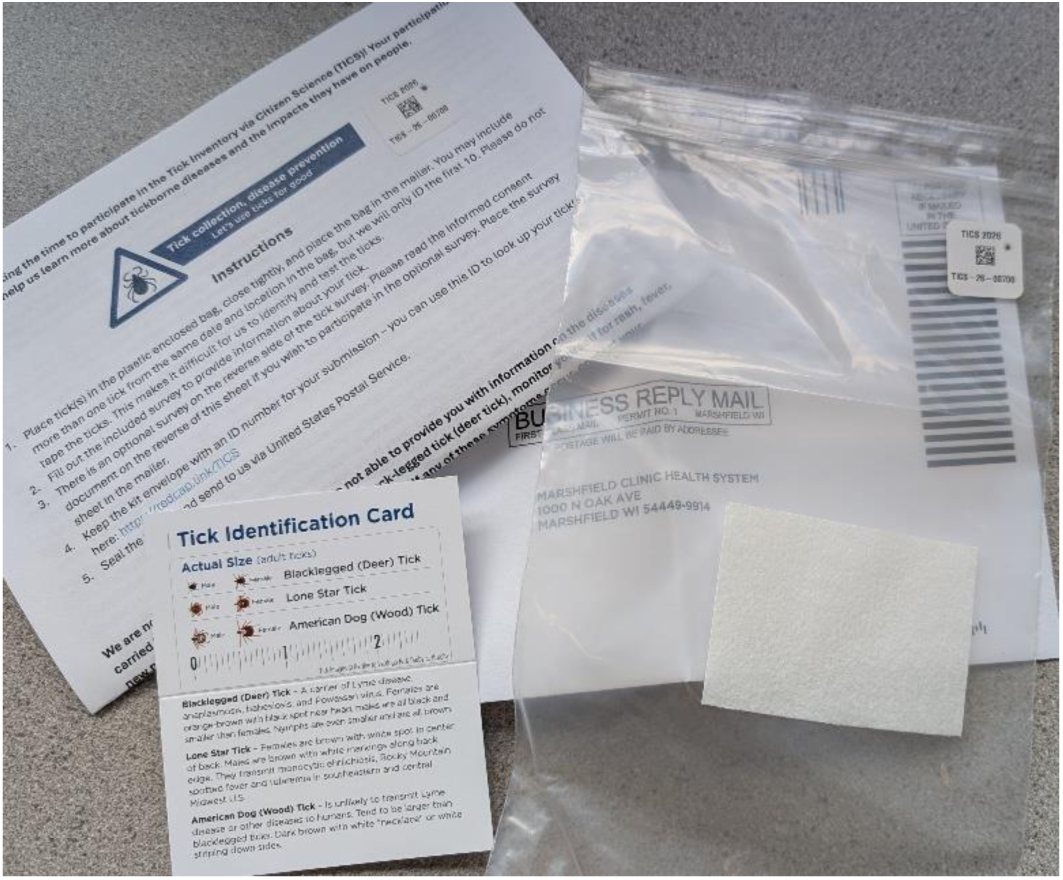
Contents of a tick collection kit.

## Methods

### Tick collection

Members of the public submitted ticks using pre-assembled collection kits. These kits included a plastic bag to contain the ticks, an information sheet, a brief survey, and a prepaid return envelope. The information sheet included a description of our study, how to safely remove a tick and when to seek medical care for a potential tickborne disease (using educational materials from the United States Centers for Disease Control and Prevention and from the Wisconsin Department of Health Services), and instructions for how to use the kit. The survey asked for information about when and where the tick was found, as well as what activity the participant was doing when they acquired the tick and what preventative measures they took. Ticks removed from animals were also accepted. Each kit was labeled with a unique identification number and barcode. The participant retained one copy of the ID on their instructions/consent form sheet so that they could look up their species identifications on our online, public dashboard (<u>redcap.link/TICS</u>). Both documents are available as Supplemental Materials.

Collection kits were distributed via countertop kiosks at community locations, handed out at local events, or mailed upon request. Community locations included clinics and hospitals in the Marshfield Clinic region of Sanford Health, veterinary clinics, visitor centers at outdoor recreation areas, coffee shops, libraries, and more. We targeted locations with high traffic locations, particularly where we thought visitors might encounter ticks, with a prominent, indoor location to display the collection kits. Kit distribution at local events was paired with tickborne disease education provided by our staff. Our scientists also appeared on local media outlets to request participation in TICS. Press releases were timed for early spring, Memorial Day, July 4^th^, Labor Day, and the start of hunting season each year to maintain awareness of the study. Members of the public used the dedicated study email address and phone number to request kits.

We also performed active surveillance via tick dragging in 2025. The locations sampled included Brunet Island State Park, Geoge Mead Wildlife Area, and Kinnickinnic State Park throughout June and July. Dragging was performed by dragging a 1m cloth along the edges of trails for a total of 750m^2^, with stops to check for ticks every 15m. Ticks found on the person dragging were also included in the total count.

The tick collection portion of this study was reviewed by our internal institutional review board (IRB) and determined not to be human subject research. Information collected via optional surveys accompanying tick submission in 2025, including demographics, prior tickborne disease diagnoses, and contact information, was approved by our IRB with a waiver of documentation of consent. The consent form included in collection kits is available in the supplemental documents.

### Tick identification

Tick collection kits were returned to our laboratory via prepaid postal mailers. Upon receipt, data from the included surveys and tick identification were collected and managed using REDCap electronic data capture tools hosted at the Marshfield Clinic Research Institute (Harris et al., 2009, 2019). Ticks were identified to the species, sex, and life stage (as possible) by trained staff. Any ticks suspected to be a species other than *I. scapularis* or *D. variabilis* were set aside for further review. Other qualities of the specimen, such as state of engorgement and whether or not its mouthparts were present, were also noted in the REDCap database. Specimens were then preserved in 70% ethanol at -20°C. Geocodes, collection dates, and species identifications were automatically uploaded to our online dashboard after entry into REDCap.

Because of the significant concern about the range expansion of *A. americanum,* participants who submitted a tick of this species in 2025 and consented to electronic contact were re-contacted to request more details about their submission. This included recent travel history and additional information about where the tick was found.

### Pathogen testing

We selected non-engorged, adult female *I. scapularis* with specific location data submitted in 2024 for pathogen testing. Prior to lysis, ticks were thawed, then washed via brief vortexing in 70% ethanol followed rinsing in nuclease-free water. Specimens were homogenized in phosphate-buffered saline via bead beating using Matrix S lysing tubes (#1169250-CF, MP Biomedical, Santa Ana, CA, USA) for two 60-second cycles at 4m/s, with a 45-second dwell between cycles. DNA was extracted from homogenized specimens using the Qiagen DNEasy Blood and Tissue Kit (#69506, Qiagen, Venlo, Netherlands). DNA concentration was measured via Qubit using the dsDNA HS assay kit (Thermo Fisher Scientific, Massachusetts, USA). DNA was eluted in 100 uL TE buffer and stored at -20C until testing.

Pathogen detection was performed using Taqman Microbial Assays Master Mix (#A50133) and primer-probe pairs specific for each pathogen were purchased from Thermo Fisher Scientific (Massachusetts, USA) (*B. burgdorferi:* #Ba07922958_s1, *B. microti:* #Pr07922949_s1, *A. phagocytophilum:* #Ba07922353_s1, *E. muris eauclairensis:* #Ba07922472_s1). Testing was performed using the QuantStudio 7 Pro System (Thermo Fisher Scientific, Massachusetts, USA), and results were determined using the Design & Analysis 2.8.0-Real-Time PCR System (2023, Thermo Fisher Scientific, Massachusetts, USA). Testing was performed following the manufacturer’s protocol, utilizing the following cycling parameters: 25°C x 2 min; 50°C x 15 min; 95°C x 2 minutes; 40 x [95°C x 3 seconds, 60°C x 1 minute]. Genomic DNA for use as positive controls for *B. burgdorferi* (#35210D-5) and *B. microti* (#NC3794365) were purchased from the American Type Culture Collection (ATCC, Viginia, USA), while genomic DNA from *A. phagocytophilum* (#ANARC) was purchased from Bio-Rad Laboratories (California, USA). A sample of genomic DNA from *E. muris eauclairensis* was provided by the CDC Rickettsial Isolate Reference Collection. Pathogen testing results were not returned to participants.

### Data analysis

Statistical analysis was performed using R, RStudio, and the R package “tidyverse” (R Core Team, 2020; RStudio Team, 2019; Wickham et al., 2019). Some calculations, such as a collection kit’s time in transit, were calculated within REDCap during data entry. Figures were generated using the R package “ggplot2” (Wickham et al., 2019). The smoothing function used over time was the conditional mean with the “loess” function (locally weighted regression). Significant differences in pathogen prevalences by geography and by month were assessed using a chi-squared test, as were significant associations between pathogens. Confidence intervals were calculated at 95% using the standard error method for proportions. Significant difference in ratios of *D. variabilis* to *I. scapularis* and adults to nymphs for active vs. passive surveillance were assessed using Fisher’s exact test.

We used a Google Maps API embedded in the REDCap data entry tool to convert addresses to geocodes. Mapping and geospatial analyses were performed using ArcGIS Pro (ESRI, Redlands, CA, USA). Shapefiles for the state of Wisconsin and its counties were obtained from the Wisconsin Department of Natural Resources Open Data Portal (https://data-wi-dnr.opendata.arcgis.com/, accessed April 22, 2026). We used Level 3 ecoregions, as defined by the United States Environmental Protection Agency (EPA), as proxies for ecosystem types. Shapefiles for ecoregions were obtained from the EPA website (https://www.epa.gov/eco-research/ecoregion-download-files-state-region-5, accessed April 22, 2026).

## Results

In the first two years of surveillance, we received 12,492 ticks. The most common tick species submitted was *D. variabilis,* followed by *I. scapularis* (Table 1). Combined, these two species comprised 99.7% of all submissions. Submissions were predominantly female (70.7% of *I. scapularis* and 54.6% of *D. variabilis*). We infrequently received larvae or nymphs of any species. The ratio of *D. variabilis* to *I. scapularis* was 2.7:1 in 2024 and 1.66:1 in 2025. We received both species from every county in Wisconsin, although the highest numbers of submissions originated in central and northern Wisconsin, particularly around Marshfield, Stevens Point, and Wausau (Figure 3).

**Figure 3.**
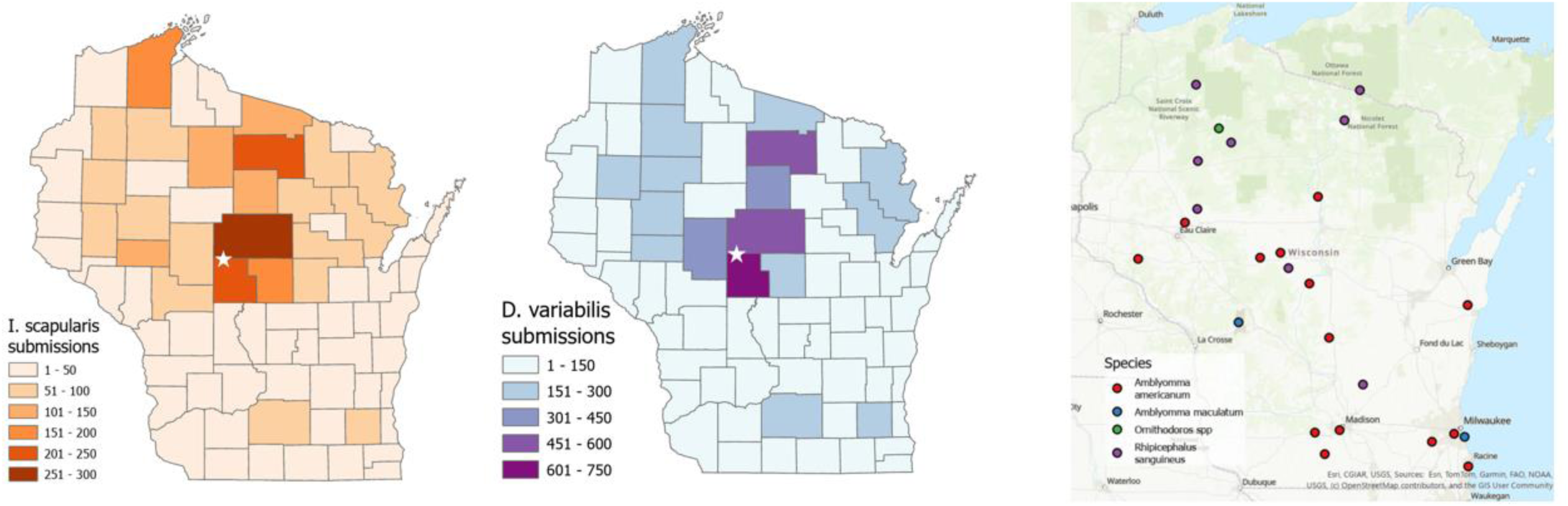
Species submitted from Wisconsin in 2024 and 2025. A) All *I. scapularis* submissions, B) All *D. variabilis* submissions, and C) other submissions. We note that the Marshfield Clinic Research Institute is located in the center of Wisconsin (white star), where we observed the highest numbers of tick submissions.

**Table 1.** Tick species identified via passive surveillance. An additional 51 submissions were unidentifiable or contained arthropods other than ticks.

|  | 2024<br>Larvae | Nymphs | Adults | 2025<br>Larvae | Nymphs | Adults |
| --- | --- | --- | --- | --- | --- | --- |
| *Ixodes scapularis* | 8 | 136 | 1534 | 14 | 179 | 2177 |
| *Dermacentor variabilis* | 0 | 5 | 4457 | 0 | 6 | 3940 |
| *Amblyomma americanum* | 0 | 0 | 13 | 0 | 3 | 6 |
| *Rhipicephalus sanguineus* | 0 | 0 | 10 | 0 | 1 | 0 |
| *Amblyomma maculatum* | 0 | 0 | 2 | 0 | 0 | 0 |
| *Ornithodoros spp.* | 0 | 0 | 0 | 0 | 0 | 1 |

We performed tick dragging at locations in central Wisconsin to learn how citizen science submissions differed from environmental sampling. We found that we were more likely to see larger ticks (both larger species and more adults) from citizen science submissions compared to tick drags. Specifically, our ratio of *D. variabilis* to *I. scapularis* was 1:5, and 64% of specimens found via tick dragging were nymphs. Both values are significantly different than those observed from citizen science submissions (p < 0.001).

Tick submissions varied seasonally across both years of sampling (Figure 4). In both 2024 and 2025, *D. variabilis* began arriving in substantial numbers in April and peaked in May, with relatively few submissions from late July through the end of tick season. In contrast, *I. scapularis* showed two annual peaks, and the timing differed each year. In 2024, *I. scapularis* submissions peaked in May, with a peak again in October. In 2025, submissions of *I. scapularis* peaked earlier, with the first peak in April and the second in October. Overall, the timing of tick season appeared relatively consistent throughout these two years of surveillance.

**Figure 4.**
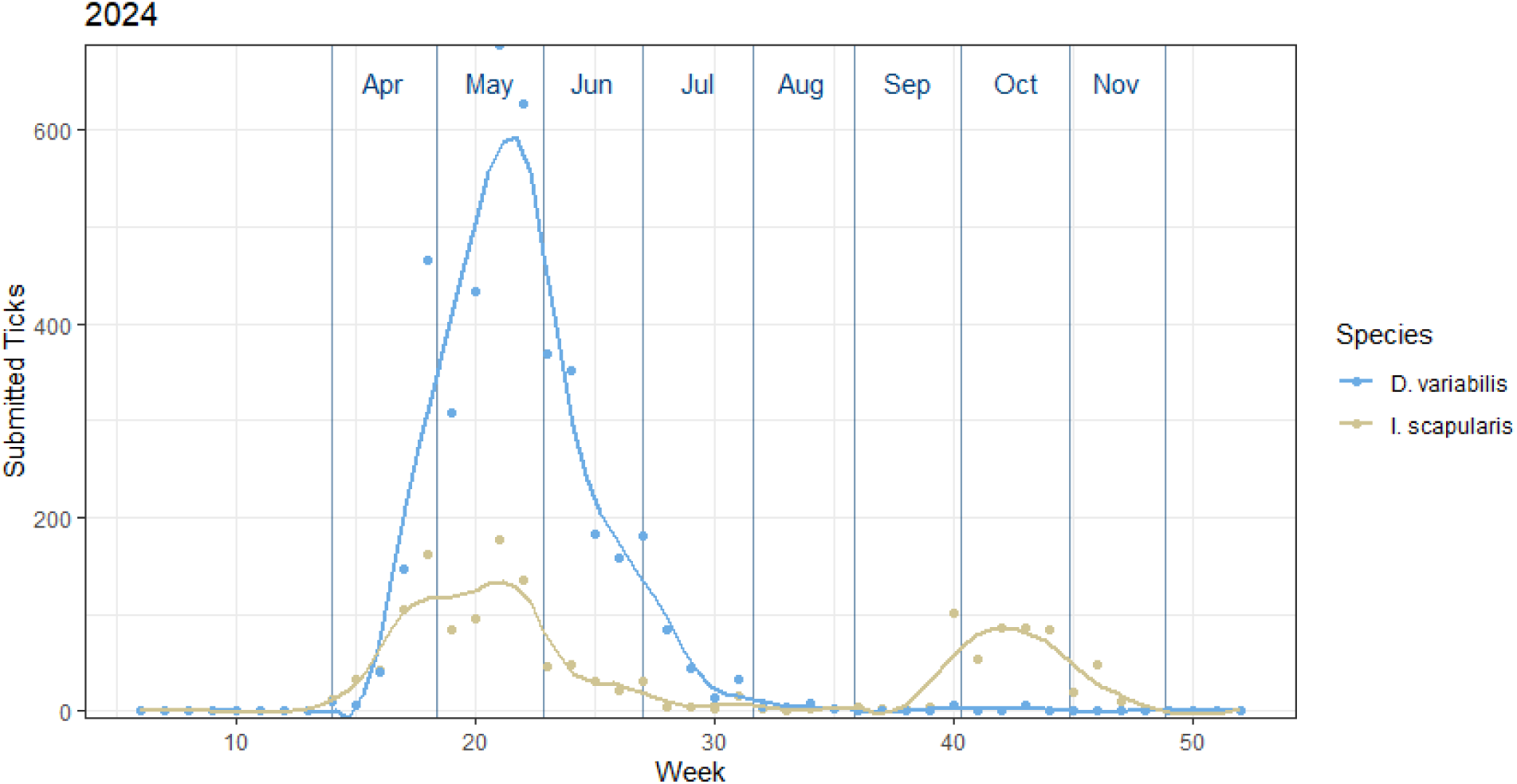
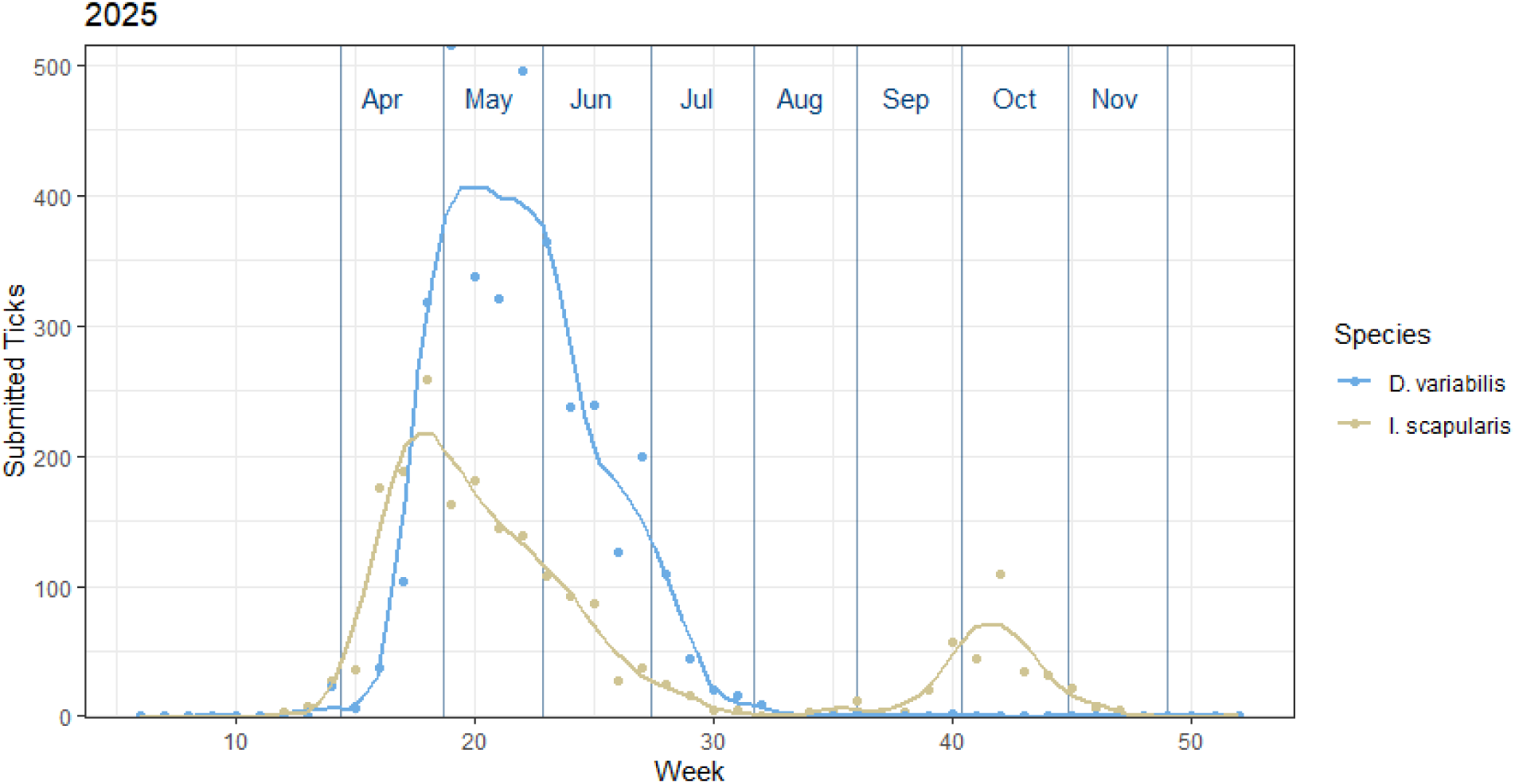
Timing of tick submissions. The timing of tick seasonality remained relatively consistent across both years for both species. However, the exception is an early April peak for *I. scapularis* in 2025 compared to its May peak in 2024. A 2^nd^ peak of submissions in October was observed in both years.

Climate and ecosystems vary across the upper Midwest, particularly from north to south. We further examined ecoregions (as defined by the EPA) at level 3 with at least 75 tick submissions for species inventory and timing of submissions. The four ecoregions analyzed were all in Wisconsin and included the Southeastern Wisconsin Till Plains, Northern Lakes and Forests, North Central Hardwood Forests, and the Driftless Area. We saw the highest numbers of submissions from the North Central Hardwood Forests and the Northern Lakes and Forests. The timing of tick season remained relatively consistent across ecosystems (Figure 5). *I. scapularis* submissions increased in the spring, then slowly dropped until mid-July. The fall peak in *I. scapularis* submissions occurred in all regions, with Northern Lakes and Forests peaking first in both years.

**Figure 5.**
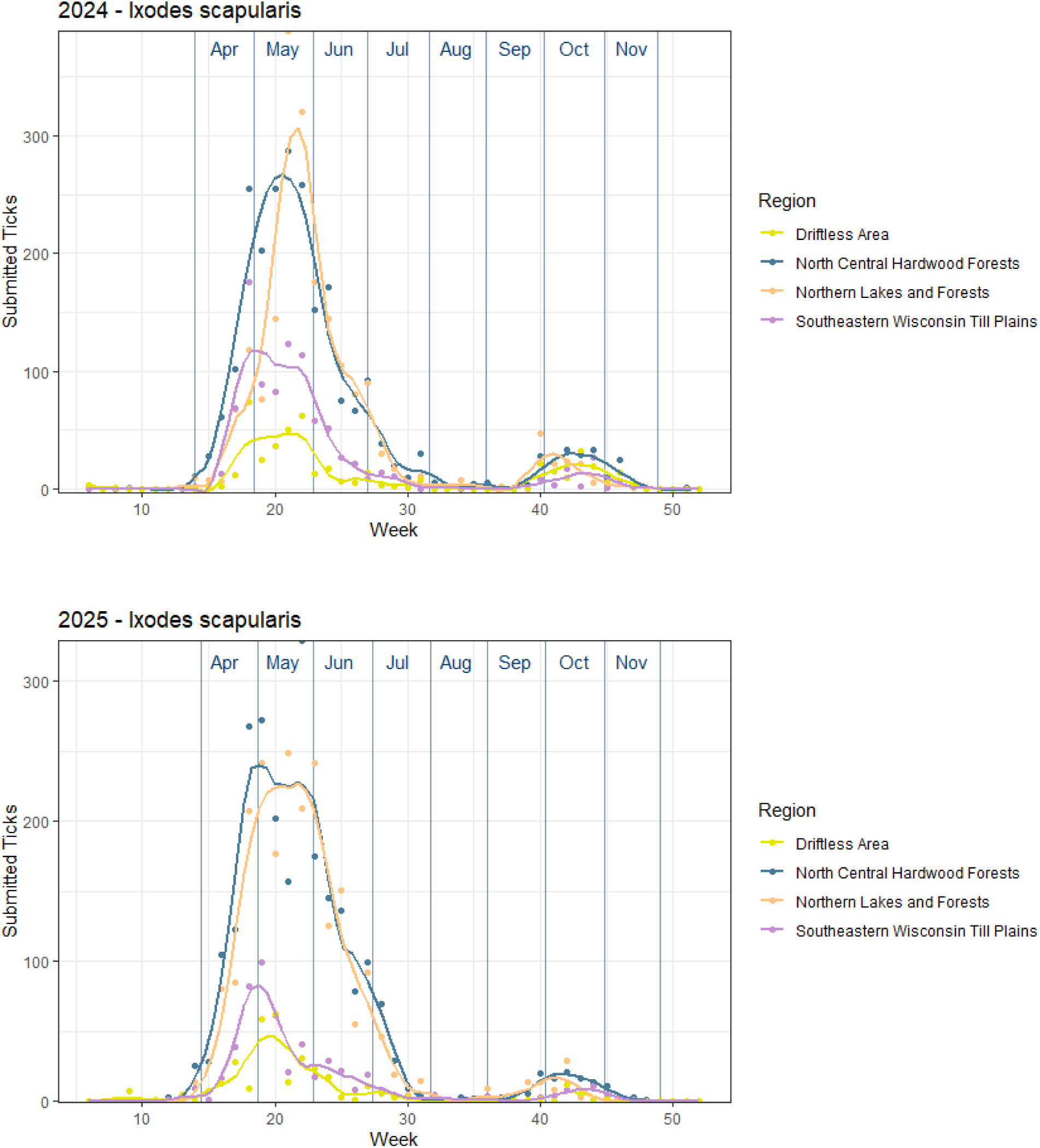
Tick seasonality by ecoregion. Timing of *I. scapularis* submissions differed by ecoregion, with our two northern ecoregions peaking sooner and higher than the southern and western regions. A fall peak was observed in all ecoregions. A map of ecoregions is available in Figure 6.

### Pathogen testing results

We tested non-engorged adult female *I. scapularis* submitted in 2024 for the four most common pathogens in our area: *B. burgdorferi, A. phagocytophilum, B. microi,* and *E. muris eauclairensis.* We tested 707 specimens in total. The overall positivity rates were 51% [95% CI: 47-55%] for *B. burgdorferi,* 9% [7-12%] for *B. microti,* 9% [7-12%] for *A. phagocytophilum,* and 3% [1-4%] for *E. muris eauclairensis. I. scapularis* can carry more than one pathogen, so we assessed associations between pathogen positivity (Table 2). Significant associations were identified between *B. burgdorferi* and *E. muris eauclairensis* and between *B. burgdorferi* and *B. microti.* Of specimens that tested positive for *B. microti,* 69% also tested positive for *B. burgdorferi* (p=0.003), and 78% of specimens positive for *E. muris eauclairensis* were also positive for *B. burgdorferi* (p = 0.038).

**Table 2.** Co-occurrence of pathogens in *I. scapularis.* An asterix indicates p < 0.05. Reported percentages are of all specimens tested. We observed 1% of specimens positive for *B. burgdorferi, A. phagocytophilum,* and *B. microti* concurrently. No specimens tested positive for all four pathogens.

|  | *B. burgdorferi* | *E. muris eauclairensis* | *A. phagocytophilum* |
| --- | --- | --- | --- |
| *E. muris eauclairensis* | 2%* |  |  |
| *A. phagocytophilum* | 6% | <1% |  |
| *B. microti* | 7%* | <1% | <1% |

We observed variation in pathogen prevalence by ecoregion (Figure 6). The highest proportion of *B. burgdorferi* positivity was found in the Driftless Area at 62% [53-73%] positive, while the lowest proportion of *B. burgdorferi* positivity, 47% [42-54%], was observed in the North Central Hardwood Forests. Positivity for *B. microti* ranged from 5% (Southeastern Wisconsin Till Plains) to 17% (Driftless Area), while positivity for *A. phagocytophilum* ranged from 4% (Driftless Area) to 15% (Northern Lakes and Forests). *E. muris eauclairensis* was not detected in any specimens from the Southeastern Wisconsin Till Plains and ranged from 1-3% in the other regions.

**Figure 6.**
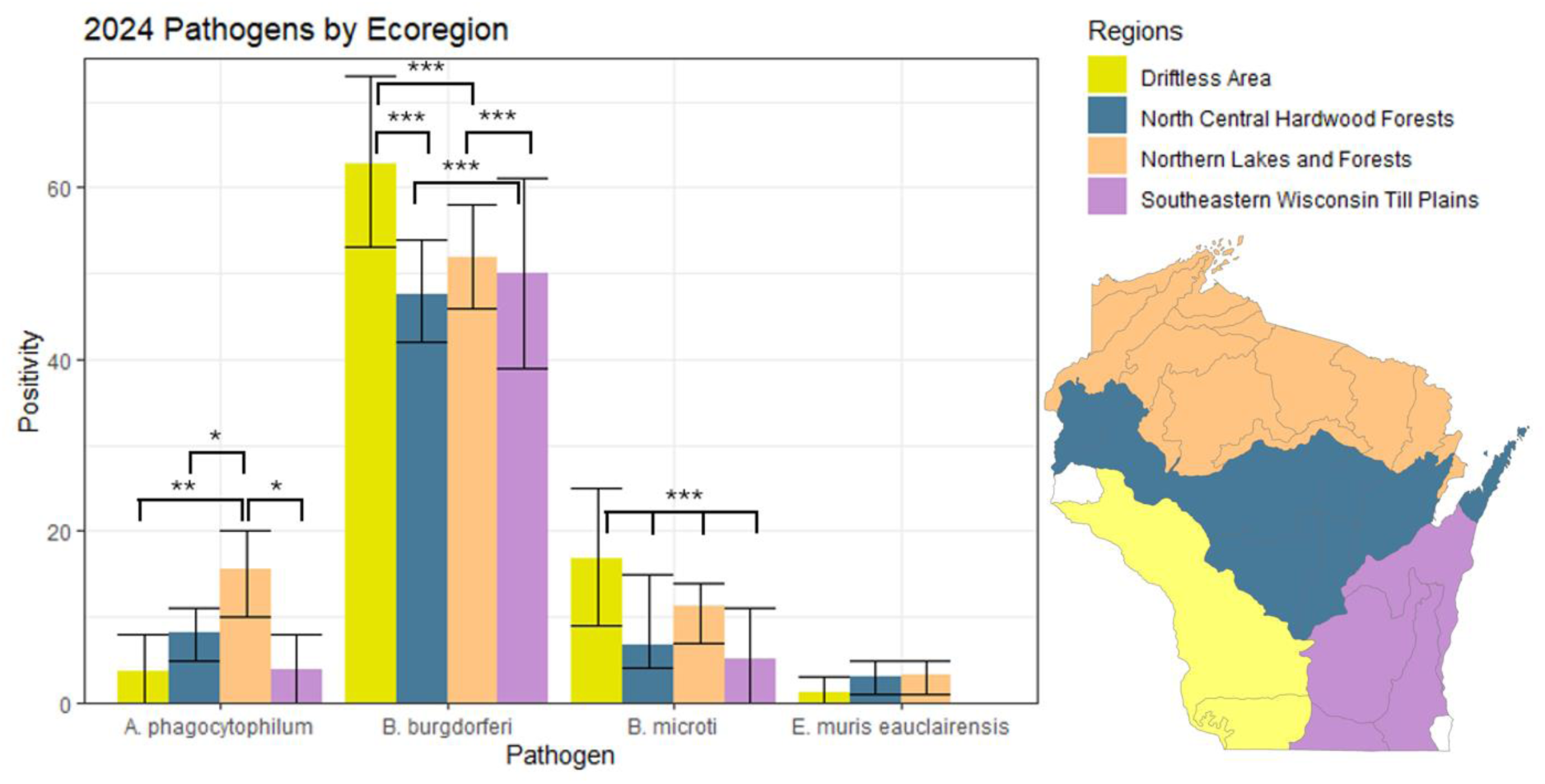
Pathogen positivity by ecoregion. We observed significant differences in pathogen prevalence between ecoregions. *B. microti* prevalence differed between all ecoregions, while *A. phagocytophilum* and *B. burgdorferi* differed between some groups. No significant differences were observed between ecoregions for *E. muris eauclairensis*.

Positivity also fluctuated throughout the year. To assess seasonality in pathogen prevalence, we analyzed each month that contained at least 10 tested specimens (Figure 7). We observed that *B. burgdorferi* positivity was higher in the fall than in the spring and summer, with percentages of positive specimens of 66% [57-75%] in October compared to 36% [27-45%] in April and 48% [40-56%] in May. All months were significantly different from each other via chi-squared test at p < 0.001 with the exceptions of April vs October (p = 0.01) and June vs November (not significant).

**Figure 7.**
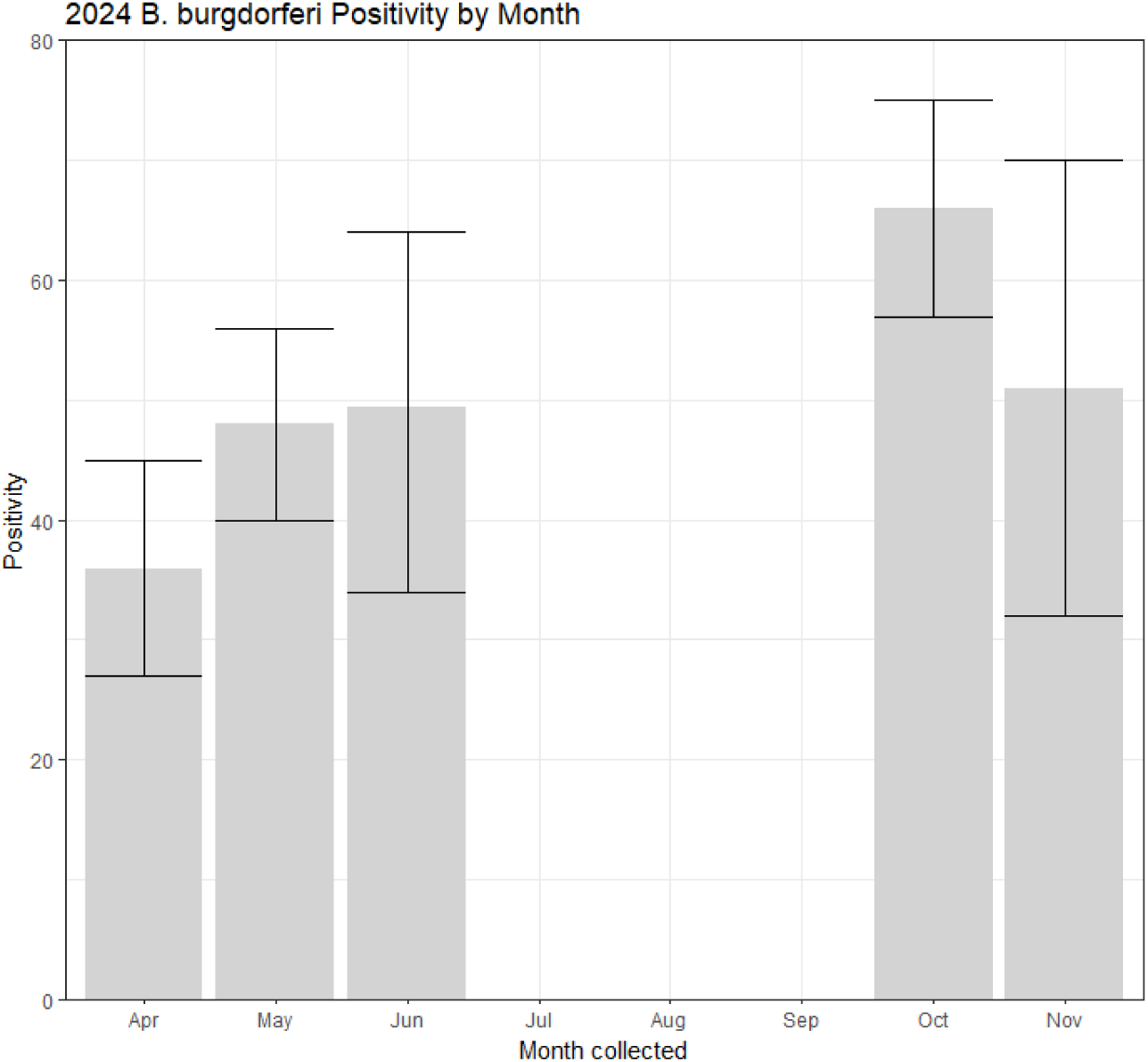
*B. burgdorferi* positivity by month. Only months with greater than 10 tested specimens are shown here; July, August, and September did not meet this threshold. All combinations except June and November are significantly different.

Lastly, we analyzed which counties in Wisconsin had positive specimens for each tested pathogen (Figure 8). *B. burgdorferi* was widespread across Wisconsin, while *A. phagocytophilum* was observed primarily in northern Wisconsin. *B. microti* had a scattered distribution across the state, while *E. muris eauclairensis* was detected rarely in sorthern Wisconsin. These results are highly dependent on sampling coverage, and positive counties are likely biased towards those with large numbers of tick submissions. The absence of a positive result in a county does not indicate the absence of the pathogen in that county, but a positive result indicates its presence. However, this data is still important because it significantly adds to the list of counties where these pathogens have been detected in *I. scapularis* (https://www.cdc.gov/ticks/data-research/facts-stats/tickborne-pathogen-surveillance-1.html, accessed April 30, 2026).

**Figure 8.**
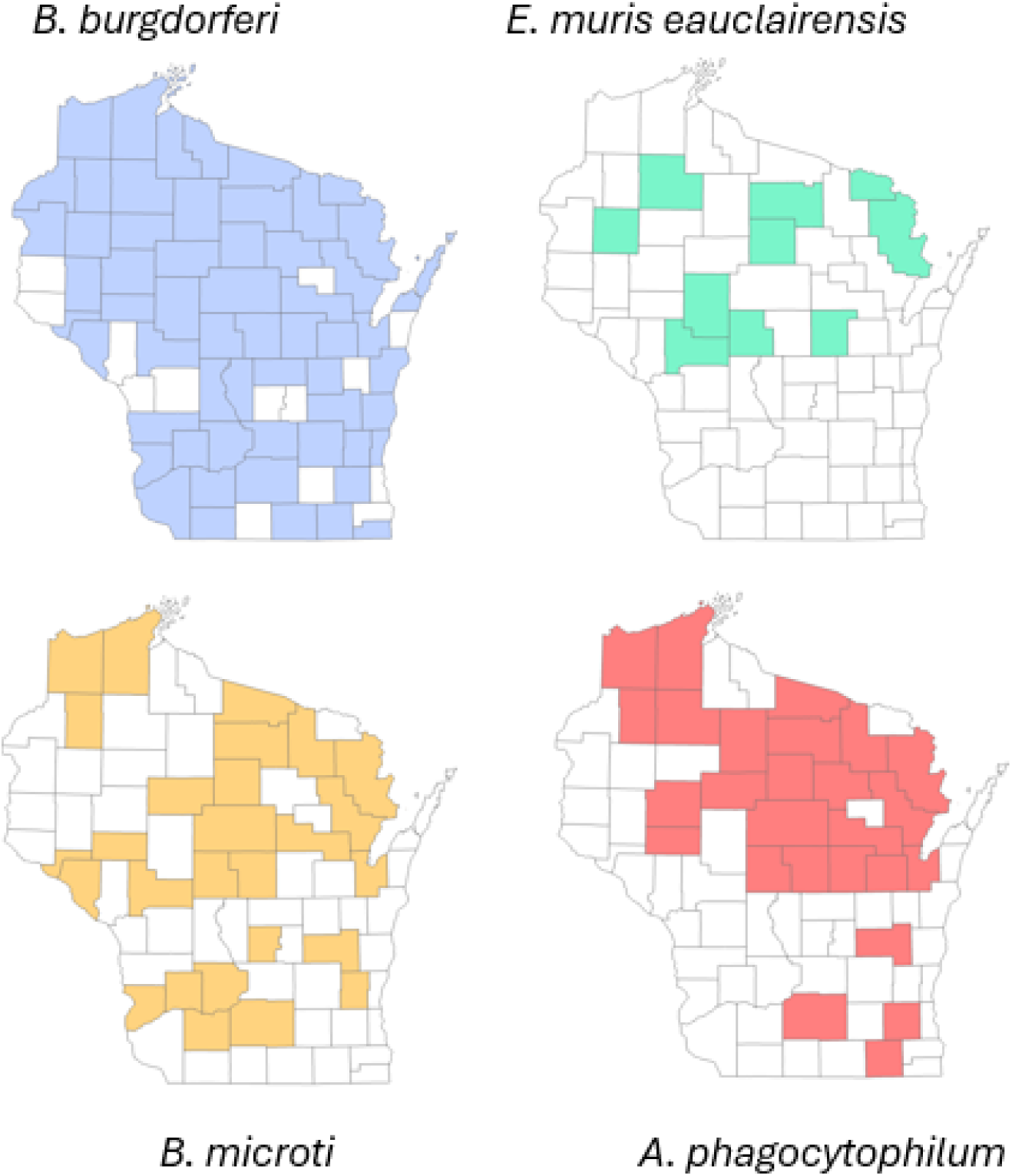
Wisconsin counties with at least one positive test result by pathogen. Counties where we found any positive specimens for each pathogen are colored, while counties with no positive specimens are white. The absence of a positive result does not indicate absence of the pathogen, as sample sizes between counties were unequal. However, these maps do expand our knowledge of where each pathogen can be found.

## Discussion

### Species Inventory

As expected, most submissions to TICS were either *I. scapularis* or *D. variabilis.* Compared to the results of tick dragging, citizen science submitted ticks tended to be larger, including a higher proportion of *D. variabilis* and of adults of any species. This is consistent with the results of other passive surveillance tick programs (Hart et al., 2022; Porter, Wachara, et al., 2021). However, it does highlight the need for increased awareness of tick bite prevention strategies in the general public, as nymphal *I. scapularis* are highly competent disease vectors. Many of our submissions came from northern and central Wisconsin. These areas both include prime tick habitat, and there is also substantial tourism from nearby metropolitan areas to the Northern Lakes and Forests ecoregion. However, we cannot rule out that proximity to our institute or a Marshfield Clinic location increased the likelihood of participation in our study.

A major concern in the upper Midwest is northward range expansion of *A. americanum.* Over two years, we received 16 specimens that we identified as *A. americanum* from Wisconsin. In our second year of collection, we were able to re-contact participants who provided an email address if they submitted an unusual specimen. This proved valuable, as it allowed us to learn that there had been recent travel to an endemic area from the participant who submitted nymphal *A. americanum.* Determination of establishment requires detection of multiple life stages of a tick. Therefore, we currently see no evidence of *A. americanum* establishment in our study area. Non-endemic ticks likely traveled on wildlife hosts to the upper Midwest, and ours is not the only study to find *A. americanum* in Wisconsin (Lee et al., 2019).

### Pathogen prevalence

Our pathogen prevalence estimates, particularly for *B. burgdorferi,* were on the higher end of previously published estimates. A study comparing prevalences of *E. muris eauclairensis* and *A. phagocytophilum* in *I. scapularis* collected either off animals or from dragging across Wisconsin from 2011-2015 found 8.9% and 5.4% positive for *A. phagocytophilum* from animals and questing ticks, respectively (Murphy et al., 2017). The values for *E. muris eauclairensis* were 2.9% positive from animal-collected ticks and 0.9% from questing ticks. A study specifically examining for *B. burgdorferi* in adult female *I. scapularis* collected by dragging in 2010-2013 found a total prevalence of 35.7%, although local prevalences in central Wisconsin and the Eau Claire region in 2013 exceeded 50% (Turtinen et al., 2015). Positivity rates of *A. phagocytophilum* in *I. scapularis* collected from hunted deer in western and east central Wisconsin in 2006 ranged between 6 and 23% (Michalski et al., 2006). Estimates of pathogen prevalences in adult *I. scapularis* collected via dragging in 2020 from Washburn County (northwestern WI) were 53% positive for *B. burgdorferi,* 3% for *B. microti,* 6% for *E. muris eauclairensis,* and 6% for *A. phagocytophilum* (Stauffer et al., 2020).

Correlations between tick infection rates and human cases of Lyme disease have been described, suggesting that these prevalence rates may serve as markers of risk to human health (Stafford et al., 1998). Specific to *B. microti,* a study in Trempealeau County in western Wisconsin found a positivity of 5% in 2013; this finding was linked to an increase in babesiosis cases seen in the Gunderson Health System, leading the authors to recommend the use of a tickborne disease diagnostic panel in this previously non-endemic area (Kowalski et al., 2015). High levels of variations between years have also been observed in tick pathogen prevalences, so testing of submitted ticks from 2025 in our study may provide different results (Foster et al., 2022).

We noted an especially high prevalence of *B. burgdorferi* in the Driftless Area ecoregion, which covers much of southwestern Wisconsin. Our estimate of 62% positivity is very high, but consistent with previous findings of higher rates in western Wisconsin compared to other areas of the state. Population genetics of *I. scapularis* indicate that two refugia for this species existed in northern and southwestern Wisconsin prior to its more recent range expansion (Dong et al., 2025). Perhaps a longer period of establishment in western vs eastern Wisconsin has led to higher infection rates with *B. burgdorferi.* Alternatively, this difference in prevalence may be driven by ecosystem traits. The Driftless Area is geologically distinct from the rest of Wisconsin, as its position on the terminal edge of glaciers resulted in bluffs and “coulees” (steep ravines). The tops of the bluffs host oak forests, which in turn support rodent populations that feed on acorns, potentially driving higher *B. burgdorferi* prevalence rates (Ostfeld et al., 2001). We must also consider bias in our study design, which depends on where people encounter ticks and who decides to participate in our study. A large number of tick submissions from the Driftless Area came from Governor Dodge and Blue Mound State Parks, so our prevalence may not be generalizable to all of western Wisconsin.

We also observed significantly higher prevalences of *B. burgdorferi* in fall compared to spring and summer. This is consistent the findings of New York’s citizen science *tick*Map effort, which attributed this phenomenon to greater numbers of adult submissions in the fall (Hart et al., 2022). However, we only tested adult females, who should all have had equal numbers of blood meals and similar to contract *B. burgdorferi* from a host. One hypothesis is that *B. burgdorferi* is less able to persist through the long winter months between feedings compared to the relatively short time frame between spring and fall. This finding highlights the continued need for tick bite prevention strategies in the fall, particularly as the timing of *I. scapularis’* fall peak coincided with hunting seasons in the upper Midwest, when many people spend increased time in wooded and brushy landscapes.

### Co-occurrence of pathogens in *I. scapularis*

We identified significant associations between *B. burgdorferi* and *B. microti,* and between *B. burgdorferi* and *E. muris eauclairensis.* Co-infections with tickborne pathogens have long been an area of concern, and our data indicates that many *I. scapularis* are carrying more than one pathogen. We have two hypotheses as to why these significant associations exist. The first is that each pathogen is responding to the same ecological drivers – for example, wildlife host diversity – that cause locally higher prevalence rates (Ratti et al., 2021). If the hosts the ticks are feeding upon are co-infected, their next host is more likely to contract more than one pathogen as well. The second hypothesis is that one pathogen facilitates entry and persistence of another in a tick. This could be through nutrient exchange or through immune evasion. For example, *B. microti* has an “arrowhead” organelle as part of its apical complex that allows it to pass through the peritrophic membrane, which separates a tick’s blood meal from its epithelial cells (Rudzinska et al., 1982). Perhaps *B. burgdorferi* takes advantage of this membrane disruption, leading to greater numbers of co-infected *I. scapularis*.

We note that co-infected *I. scapularis* were not rare, with 14% of tested specimens containing more than one pathogen. *B. burgdorferi* was by far the most common pathogen we detected, consistent with Lyme disease as the most common tickborne disease in Wisconsin (Rau et al., 2020). Lyme serology testing is often not positive in early disease, so clinical signs and symptoms may be helpful in distinguishing Lyme from other tickborne diseases, the latter of which can be screened for with other diagnostic tests, including a PCR panel. Anaplasmosis and ehrlichiosis are often characterized by rapid onset fever, chills, headache, and malaise, along with low white blood cell and platelet counts and elevated liver enzyme levels (Ismail & McBride, 2017) (https://www.cdc.gov/anaplasmosis/hcp/clinical-signs/, https://www.cdc.gov/ehrlichiosis/hcp/clinical-signs/, accessed May 6, 2026). Symptom onset with babesiosis is usually delayed, and some patients may be asymptomatic; others may present with fever, chills, headache, and body aches, and platelet counts are often low. Older people and those with co-morbidities or weakened immune systems are at increased risk for serious disease, including hemolytic anemia (“Babesiosis,” 2020; Rajapakse & Bakirhan, 2021) (https://www.cdc.gov/babesiosis/signs-symptoms/index.html, accessed May 6, 2026). While *E. muris eauclairensis* was the least common pathogen detected in our study and is endemic to the upper Midwest, travel history to the southern United States, where *E. chaffeensis* is endemic, may increase exposure risk to ehrlichiosis. Fortunately, doxycycline is an effective treatment for Lyme disease, ehrlichiosis, and anaplasmosis. Babesiosis, however, is not responsive to doxycycline, instead requiring a combination of atovaquone and azithromycin for treatment (Krause et al., 2021).

### Continuing efforts

TICS is ongoing, and year three of tick collection began in March of 2026. We are continuing to monitor non-endemic tick species in collaboration with public health departments. We preserve all ticks for future testing, so that we can continue measuring pathogen prevalences in subsequent years as resources permit. We are also currently studying the microbiomes of ticks from 2024 submissions. We are using 16S amplicon sequencing to compare microbiome composition between *I. scapularis* with and without *B. burgdorferi* in the hopes of identifying non-pathogenic bacteria that either foster or inhibit a tick’s ability to carry a pathogen. We are also piloting 16S sequencing as a disease-agnostic surveillance technique on *D. variabilis*.

The surveys included in each kit provide a wealth of information about people’s knowledge, attitudes, and practices surrounding tickborne diseases, along with a paired tick encounter and the tick itself. We are continuing to analyze this data and to propose new survey types to include with tick submission. This year, we have launched an initiative called the Lyme Experiences and Narratives Study, which collects long-form, open-ended stories about people’s experiences with tickborne diseases (redcap.link/TICSLENS). We started this study after talking to participants and realizing that there are nuance and breadth regarding this topic that cannot be captured in multiple choice surveys.

Because we are part of a health system, we can also link the ecology of ticks and their pathogens to what we see in our clinics. By comparing case counts and test results for tickborne diseases to our local prevalence estimates, we can learn how closely the two match. This approach may provide new insights into risk assessment for tickborne diseases, both for individuals seeking to reduce their risk and for clinicians trying to diagnose their patients.

We will continue using this opportunity to provide high quality tickborne disease information to the public. We are currently developing a methods paper to help other researchers develop their own citizen science programs, particularly in rural areas. Our goal is to increase people’s trust and confidence in the results through participation in citizen science initiatives (Bedessem et al., 2021). This feels critically important for tickborne diseases, where there is much that the scientific community cannot explain and misinformation is abundant. The combined efforts of citizen scientists as described here may assist in more accurate risk assessment to aid in the reduction of tickborne disease burden across the Midwest.

## Acknowledgements

Funding for this project was provided through philanthropic support of Marshfield Clinic Research Institute from the Frank and Betty Koller Trusts and the Sanford Health Foundation – Marshfield.

## References

Babesiosis. (2020). In *Hunter’s Tropical Medicine and Emerging Infectious Diseases* (pp. 799–802). Elsevier. 10.1016/B978-0-323-55512-8.00105-8

Bedessem, B., Gawrońska-Nowak, B., & Lis, P. (2021). Can citizen science increase trust in research? A case study of delineating Polish metropolitan areas. Journal of Contemporary European Research, 17(2). 10.30950/jcer.v17i2.1185

Corrin, T., Greig, J., Harding, S., Young, I., Mascarenhas, M., & Waddell, L. A. (2018). Powassan virus, a scoping review of the global evidence. Zoonoses and Public Health, 65(6), 595–624. 10.1111/zph.12485

Diuk-Wasser, M. A., VanAcker, M. C., & Fernandez, M. P. (2021). Impact of Land Use Changes and Habitat Fragmentation on the Eco-epidemiology of Tick-Borne Diseases. Journal of Medical Entomology, 58(4), 1546–1564. 10.1093/jme/tjaa209

Dong, D., Paskewitz, S. M., Tsao, J. I., & Schoville, S. D. (2025). Genetic and Landscape Connectivity of Blacklegged Ticks During Range Expansion in Select States of the Midwestern USA. Ecology and Evolution, 15(10), e72360. 10.1002/ece3.72360

Eilbert, W., & Matella, A. (2024). Tick-Borne Diseases. Emergency Medicine Clinics, 42(2), 287–302. 10.1016/j.emc.2024.01.004

Eisen, L., & Eisen, R. J. (2021). Benefits and Drawbacks of Citizen Science to Complement Traditional Data Gathering Approaches for Medically Important Hard Ticks (Acari: Ixodidae) in the United States. Journal of Medical Entomology, 58(1), 1–9. 10.1093/jme/tjaa165

Foster, E., Burtis, J., Sidge, J. L., Tsao, J. I., Bjork, J., Liu, G., Neitzel, D. F., Lee, X., Paskewitz, S., Caporale, D., & Eisen, R. J. (2022). Inter-annual variation in prevalence of *Borrelia burgdorferi* sensu stricto and *Anaplasma phagocytophilum* in host-seeking *Ixodes scapularis* (Acari: Ixodidae) at long-term surveillance sites in the upper midwestern United States: Implications for public health practice. Ticks and Tick-Borne Diseases, 13(2), 101886. 10.1016/j.ttbdis.2021.101886

Gould, L. H., Fee, R., White, J., Webb, N., Carlyle, M., Dick, L., Tan, Y., Walker, V., Angulo, F. J., Moïsi, J. C., Stark, J. H., & Pugh, S. (2025). Risk factors for Lyme disease among residents of rural, suburban, and urban areas in the United States: A case-control study. American Journal of Epidemiology, 194(8), 2287–2294. 10.1093/aje/kwae368

Harris, P. A., Taylor, R., Minor, B. L., Elliott, V., Fernandez, M., O’Neal, L., McLeod, L., Delacqua, G., Delacqua, F., Kirby, J., & Duda, S. N. (2019). The REDCap consortium: Building an international community of software platform partners. Journal of Biomedical Informatics, 95, 103208. 10.1016/j.jbi.2019.103208

Harris, P. A., Taylor, R., Thielke, R., Payne, J., Gonzalez, N., & Conde, J. G. (2009). Research electronic data capture (REDCap)—A metadata-driven methodology and workflow process for providing translational research informatics support. Journal of Biomedical Informatics, 42(2), 377–381. 10.1016/j.jbi.2008.08.010

Hart, C. E., Bhaskar, J. R., Reynolds, E., Hermance, M., Earl, M., Mahoney, M., Martinez, A., Petzlova, I., Esterly, A. T., & Thangamani, S. (2022). Community engaged tick surveillance and tickMAP as a public health tool to track the emergence of ticks and tick-borne diseases in New York. PLOS Global Public Health, 2(6), e0000215. 10.1371/journal.pgph.0000215

Hecht, J. A., Allerdice, M. E. J., Dykstra, E. A., Mastel, L., Eisen, R. J., Johnson, T. L., Gaff, H. D., Varela-Stokes, A. S., Goddard, J., Pagac, B. B., Paddock, C. D., & Karpathy, S. E. (2019). Multistate Survey of American Dog Ticks (Dermacentor variabilis) for Rickettsia Species. Vector-Borne and Zoonotic Diseases, 19(9), 652–657. 10.1089/vbz.2018.2415

Ismail, N., & McBride, J. W. (2017). Tick-Borne Emerging Infections: Ehrlichiosis and Anaplasmosis. Clinics in Laboratory Medicine, 37(2), 317–340. 10.1016/j.cll.2017.01.006

Kowalski, T. J., Jobe, D. A., Dolan, E. C., Kessler, A., Lovrich, S. D., & Callister, S. M. (2015). The Emergence of Clinically Relevant Babesiosis in Southwestern Wisconsin. 114(4).

Krause, P. J., Auwaerter, P. G., Bannuru, R. R., Branda, J. A., Falck-Ytter, Y. T., Lantos, P. M., Lavergne, V., Meissner, H. C., Osani, M. C., Rips, J. G., Sood, S. K., Vannier, E., Vaysbrot, E. E., & Wormser, G. P. (2021). Clinical Practice Guidelines by the Infectious Diseases Society of America (IDSA): 2020 Guideline on Diagnosis and Management of Babesiosis. Clinical Infectious Diseases, 72(2), e49–e64. 10.1093/cid/ciaa1216

Kugeler, K. J., Schwartz, A. M., Delorey, M. J., Mead, P. S., & Hinckley, A. F. (2021). Estimating the Frequency of Lyme Disease Diagnoses, United States, 2010–2018. Emerging Infectious Diseases, 27(2), 616–619. 10.3201/eid2702.202731

Lee, X., Murphy, D. S., Johnson, D. H., & Paskewitz, S. M. (2019). Passive Animal Surveillance to Identify Ticks in Wisconsin, 2011–2017. Insects, 10(9). 10.3390/insects10090289

Mead, P., Hinckley, A., & Kugeler, K. (2024). Lyme Disease Surveillance and Epidemiology in the United States: A Historical Perspective. The Journal of Infectious Diseases, 230(Supplement_1), S11–S17. 10.1093/infdis/jiae230

Michalski, M., Rosenfield, C., Erickson, M., Selle, R., Bates, K., Essar, D., & Massung, R. (2006). Anaplasma phagocytophilum in central and western Wisconsin: A molecular survey. Parasitology Research, 99(6), 694–699. 10.1007/s00436-006-0217-9

Murphy, D. S., Lee, X., Larson, S. R., Johnson, D. K. H., Loo, T., & Paskewitz, S. M. (2017). Prevalence and Distribution of Human and Tick Infections with the Ehrlichia muris-Like Agent and Anaplasma phagocytophilum in Wisconsin, 2009–2015. Vector-Borne and Zoonotic Diseases, 17(4), 229–236. 10.1089/vbz.2016.2055

Ostfeld, R. S., Schauber, E. M., Canham, C. D., Keesing, F., Jones, C. G., & Wolff, J. O. (2001). Effects of Acorn Production and Mouse Abundance on Abundance and Borrelia burgdorferi Infection Prevalence of Nymphal Ixodes scapularis Ticks. Vector-Borne and Zoonotic Diseases, 1(1), 55–63. 10.1089/153036601750137688

Porter, W. T., Barrand, Z. A., Wachara, J., DaVall, K., Mihaljevic, J. R., Pearson, T., Salkeld, D. J., & Nieto, N. C. (2021). Predicting the current and future distribution of the western black-legged tick, Ixodes pacificus, across the Western US using citizen science collections. PLOS ONE, 16(1), e0244754. 10.1371/journal.pone.0244754

Porter, W. T., Wachara, J., Barrand, Z. A., Nieto, N. C., & Salkeld, D. J. (2021). Citizen Science Provides an Efficient Method for Broad-Scale Tick-Borne Pathogen Surveillance of Ixodes pacificus and Ixodes scapularis across the United States. MSphere, 6(5), 10.1128/msphere.00682-21. 10.1128/msphere.00682-21

Pritt, B. S., Allerdice, M. E. J., Sloan, L. M., Paddock, C. D., Munderloh, U. G., Rikihisa, Y., Tajima, T., Paskewitz, S. M., Neitzel, D. F., Hoang Johnson, D. K., Schiffman, E., Davis, J. P., Goldsmith, C. S., Nelson, C. M., & Karpathy, S. E. (2017). Proposal to reclassify Ehrlichia muris as Ehrlichia muris subsp. Muris subsp. Nov. And description of Ehrlichia muris subsp. Eauclairensis subsp. Nov., a newly recognized tick-borne pathogen of humans. International Journal of Systematic and Evolutionary Microbiology, 67(7), 2121–2126. 10.1099/ijsem.0.001896

R Core Team. (2020). R: A language and environment for statistical computing. R Foundation for Statistical Computing. https://www.R-project.org/

Rajapakse, P., & Bakirhan, K. (2021). Autoimmune Hemolytic Anemia Associated With Human Babesiosis. Journal of Hematology, 10(2), 41–45. 10.14740/jh820

Ratti, V., Winter, J. M., & Wallace, D. I. (2021). Dilution and amplification effects in Lyme disease: Modeling the effects of reservoir-incompetent hosts on Borrelia burgdorferi sensu stricto transmission. Ticks and Tick-Borne Diseases, 12(4), 101724. 10.1016/j.ttbdis.2021.101724

Rau, A., Munoz-Zanzi, C., Schotthoefer, A. M., Oliver, J. D., & Berman, J. D. (2020). Spatio-Temporal Dynamics of Tick-Borne Diseases in North-Central Wisconsin from 2000–2016. International Journal of Environmental Research and Public Health, 17(14). 10.3390/ijerph17145105

Ripoche, M., Gasmi, S., Adam-Poupart, A., Koffi, J. K., Lindsay, L. R., Ludwig, A., Milord, F., Ogden, N. H., Thivierge, K., & Leighton, P. A. (2018). Passive Tick Surveillance Provides an Accurate Early Signal of Emerging Lyme Disease Risk and Human Cases in Southern Canada. Journal of Medical Entomology, 55(4), 1016–1026. 10.1093/jme/tjy030

Riva, H. R., Nguyen, V. K., & Cervantes, J. L. (2023). STARI: The Other Lyme Disease? Clinical Infection and Immunity, 8(1), 5–12. 10.14740/cii.v8i1.164

Rodino, K. G., & Pritt, B. S. (2022). When to Think About Other Borreliae: Hard Tick Relapsing Fever (Borrelia miyamotoi), Borrelia mayonii, and Beyond. Infectious Disease Clinics, 36(3), 689–701. 10.1016/j.idc.2022.04.002

Rosenberg, R., Lindsey, N. P., Fischer, M., Gregory, C. J., Hinckley, A. F., Mead, P. S., Paz-Bailey, G., Waterman, S. H., Drexler, N. A., Kersh, G. J., Hooks, H., Partridge, S. K., Visser, S. N., Beard, C. B., & Petersen, L. R. (2018). Vital Signs: Trends in Reported Vectorborne Disease Cases - United States and Territories, 2004-2016. MMWR. Morbidity and Mortality Weekly Report, 67(17), 496–501. 10.15585/mmwr.mm6717e1

RStudio Team. (2019). *RStudio: Integrated Development for R*. RStudio, Inc. http://www.rstudio.com/

Rudzinska, M. A., Spielman, A., Lewengrub, S., Piesman, J., & Karakashian, S. (1982). Penetration of the peritrophic membrane of the tick by Babesia microti. Cell and Tissue Research, 221(3), 471–481. 10.1007/BF00215696

Sonenshine, D. E. (2018). Range Expansion of Tick Disease Vectors in North America: Implications for Spread of Tick-Borne Disease. International Journal of Environmental Research and Public Health, 15(3), 3. 10.3390/ijerph15030478

Stafford, K. C., Cartter, M. L., Magnarelli, L. A., Ertel, S. H., & Mshar, P. A. (1998). Temporal correlations between tick abundance and prevalence of ticks infected with Borrelia burgdorferi and increasing incidence of Lyme disease. Journal of Clinical Microbiology, 36(5), 1240–1244. 10.1128/JCM.36.5.1240-1244.1998

Stauffer, M. T., Mandli, J., Pritt, B. S., Stauffer, W., Sloan, L. M., Zembsch, T., Lee, X., & Paskewitz, S. M. (2020). Detection of zoonotic human pathogens in Ixodes scapularis in Wisconsin-2017. Journal of Vector Ecology : Journal of the Society for Vector Ecology, 45(1), 147–149. 10.1111/jvec.12384

Turtinen, L. W., Kruger, A. N., & Hacker, M. M. (2015). Prevalence of Borrelia burgdorferi in adult female ticks (Ixodes scapularis), Wisconsin 2010–2013. Journal of Vector Ecology, 40(1), 195–197. 10.1111/jvec.12152

Wickham, H., Averick, M., Bryan, J., Chang, W., McGowan, L. D., François, R., Grolemund, G., Hayes, A., Henry, L., Hester, J., Kuhn, M., Pedersen, T. L., Miller, E., Bache, S. M., Müller, K., Ooms, J., Robinson, D., Seidel, D. P., Spinu, V., … Yutani, H. (2019). Welcome to the Tidyverse. Journal of Open Source Software, 4(43), 1686. 10.21105/joss.01686

