## Supplemental Materials for "Ticks and tickborne diseases in the upper Midwestern United States: role for citizen science in assessing exposure risk"

**Thank you for taking the time to participate in the Tick Inventory via Citizen Science (TICS)! Your participation will help us learn more about tickborne diseases and the impacts they have on people.**

Tick collection, disease prevention
Let's use ticks for good

### Instructions

1. Place tick(s) in the plastic enclosed bag, close tightly, and place the bag in the mailer. You may include more than one tick from the same date and location in the bag, but we will only ID the first 10. Please do not tape the ticks. This makes it difficult for us to identify and test the ticks.
2. Fill out the included survey to provide information about your tick.
3. There is an optional survey on the reverse side of the tick survey. Please read the informed consent document on the reverse of this sheet if you wish to participate in the optional survey. Place the survey sheet in the mailer.
4. Keep the kit envelope with an ID number for your submission – you can use this ID to look up your tick(s) here: https://redcap.link/TICS
5. Seal the mailer and send to us via United States Postal Service.

**We are not a diagnostic laboratory, and we are not able to provide you with information on the diseases carried by your tick. If you have been bitten by a black-legged tick (deer tick), monitor yourself for rash, fever, new muscle and joint pain, and new fatigue for 30 days. If any of these symptoms occur, contact your healthcare provider immediately.**

How to remove an attached tick (from the US Centers for Disease Control):

1. Use clean, fine-tipped tweezers to grasp the tick as close to the skin's surface as possible.

2. Pull upward with steady, even pressure. Don't twist or jerk the tick; this can cause the mouth-parts to break off and remain in the skin. If this happens, remove the mouth-parts with tweezers. If you cannot remove the mouth easily with tweezers, leave it alone and let the skin heal.

3. After removing the tick, thoroughly clean the bite area and your hands with rubbing alcohol or soap and water.

4. Never crush a tick with your fingers. Dispose of a live tick by putting it in alcohol, placing it in a sealed bag/container, wrapping it tightly in tape, or flushing it down the toilet.

#### Information about your Experiences with Tickborne Disease: Informed Consent Document

**This document provides you with the necessary information to consent to the survey titled "Information about your experiences with tickborne disease." Only fill out this survey if you are over 18 years of age. By sending in this survey, you consent to being a part of this research study. You do not need to consent or be an adult to send us your tick and fill out the survey "Information about your tick(s)." Please keep this document for your records. If you have any questions about this document, contact us at 715-221-6462.**

**Purpose:** The purpose of the survey "Information about your experiences with tickborne disease" is to learn more about who is exposed to tickborne disease and how that exposure occurred. This study consists of the included survey and one optional follow-up survey via email. This study is being conducted by Jennifer Meece, PhD & Alexandra Linz, PhD, at the Marshfield Clinic Research Institute (MCRI). For more information about this research, you may contact either investigator at 715-221-6462.

**How to participate:** You are invited to participate in this research if you are at least 18 years of age and submitting a tick. Taking part in this research is voluntary. You can choose not to be in this study or stop at any time. If you are an adult, read this consent form and then fill out the survey. Returning the survey is your consent to the research study. If you agree to provide us your contact information, we email you an additional survey at the end of the tick season. You are not required to provide your contact information when filling out the survey. Not filling out this survey will not impact the results of your tick submission.

**Risks & Benefits:** There are no major risks from being in this study. As with all research, there is a chance that confidentiality could be compromised. Extensive security and confidentiality procedures are used to decrease the chance of this happening. Being in this study will not help you directly. This study will help us learn about who is at risk of contracting tickborne diseases. This may help others in the future.

**Confidentiality:** The information we are collecting in this survey is your protected health information. Your paper survey will be reviewed and recorded by trained research staff only. Electronic data will be stored securely by MCRI. Your information collected as part of this research will not be distributed or used for future research studies without your consent, even if your identifiers are removed. Summarized, de-identified data from this study may be published in a scientific journal.

**Electronic Communication:** If you provide your email address for the optional follow-up to this survey, you agree that researchers at MCRI may use email to communicate with you regarding this research study. Electronic communication is not secure when sent, stored, or viewed on a personal device. By sending us the survey with your email address on it, you accept all risk of loss of privacy or confidentiality associated with the use of electronic communication for this research study. You may withdraw your consent for electronic communication by notifying us at any time.

**Rights of Research Subjects:** Being in this study is voluntary. Refusing to participate or discontinuing participation at any time will involve no penalty or loss of benefits to which you are otherwise entitled. If you choose not to be part of this research, your relationship with your doctor and this institution will not change. You are not giving up any legal rights by signing this consent document and taking part in this research study. If you have any questions about your rights as a research subject, you may contact the MCRI's Institutional Review Board (IRB) at 1-800-782-8581 ext. 9-3022. The IRB is responsible for helping protect rights and welfare of human research subjects. You may also call this number to discuss problems and concerns, to request information and ask questions, and to offer input.

**Authorization to Use or Disclosed Protected Health Information for Research:** Researchers at MCRI are required by HIPAA, the federal privacy law, and other state privacy laws, to get permission to use and/or release identifiable health information from you for research purposes. If you agree to take part in this research study, you agree to provide permission to use and/or release your identifiable health information collected in our surveys for research purposes.

**What personal health information will be used or disclosed and for what purpose?** The types of identifiable information that may be used by MCRI staff include your demographics, medical diagnoses, and email address. We will use this information to contact you with a follow-up survey and to determine who is being diagnosed with tickborne diseases. Your information may be shared with individuals and groups at MCRI responsible for upholding research regulations. Your data will not be shared with individuals or groups external to MCRI.

**How long will my authorization last, and can I change my mind?** Your authorization to use and/or disclose your health information for this research does not have an end date. You may take back your authorization at any time but will have to do so in writing. Your cancelled authorization will not apply to information that was gathered before you took back your authorization. If you take back your authorization, you can no longer take part in the above-referenced research. To take back your authorization, write a note stating this decision and send it to: Jennifer Meece, Integrated Research and Development Laboratory, 1000 N. Oak Ave, Marshfield, WI 54449.

### Information about your tick(s)

Please provide information on the included survey sheet about where and when you found your tick. If you have multiple ticks in the collection tube from the same date and location, please fill out the survey only once.

Where did you pick up this tick?  Please provide **GPS coordinates or an address** if possible:

________________________________________

________________________________________

________________________________________

What date did you pick up this tick? ___________

Was this tick embedded in skin? *This means its mouthparts were embedded in skin – even a non-attached tick can grip tightly and be difficult to remove. Please see the instruction sheet for how to remove an attached tick.* ***Do not remove the tick using tape as its mouthparts are more likely to be retained under the skin.***

- ☐ Yes
- ☐ No

What species was this tick found on?

- ☐ Human
- ☐ Dog
- ☐ Cat
- ☐ Horse
- ☐ Other: ______________________

BARCODE

What activity were you doing when you picked up this tick?

- ☐ Hiking/walking
- ☐ Hunting
- ☐ Yardwork
- ☐ Working – what is your occupation?

  ________________________________
- ☐ Other: ___________________

What tick precautions did you take before or after you picked up this tick?

- ☐ Tick check
- ☐ DEET
- ☐ Permethrin-treated clothing (including Insect Shield®, InsectGuard®)
- ☐ Long sleeves/pants
- ☐ Shower after activity
- ☐ Avoid tall grass and brush
- ☐ Other:____________________

**See reverse for an optional survey about your experiences with tickborne disease.**

MARSHFIELD CLINIC RESEARCH INSTITUTE IN COLLABORATION WITH SANFORD HEALTH

### Information about your experiences with tickborne disease (optional)

Please make sure to fill out the survey on the reverse side with information about your tick, and only fill out this survey if your tick was found on yourself and you are at least 18 years old. Make sure you read the Informed Consent and HIPAA Authorization Document on the back of the instruction sheet before you take the survey. Please keep the instruction sheet and consent document.

**By returning this survey, you consent to being a part of the research study described in the included Informed Consent and HIPAA Authorization document. Only people who are at least 18 years old may participate in this survey.**

What is your age?

- ☐ 18-33
- ☐ 34-48
- ☐ 49-64
- ☐ 65-78
- ☐ 79+

Are you:

- ☐ Male
- ☐ Female
- ☐ Prefer not to say

Have you ever been diagnosed with any of the following tickborne diseases?

- ☐ Lyme disease
- ☐ Ehrlichiosis
- ☐ Babesiosis
- ☐ Anaplasmosis
- ☐ Rocky Mountain Spotted Fever
- ☐ Alpha-gal syndrome
- ☐ Other _______________

Would you be willing to be contacted by us at the end of the tick season to learn if you contracted a tickborne diseasethis year? This survey will only be via email. Please print clearly to ensure that we are able to email you. *We may also use your email address to contact you for more information if you submit an unusual tick.*

- ☐ Yes – please provide your email address:

If your email address uses one of these common domains, please circle it instead of writing it above:

@aol.com @charter.net @gmail.com @hotmail.com

@tds.net @outlook.com @yahoo.com

- ☐ No

Would you be willing to contact us if you are diagnosed with a tickborne infection prior to us following up?

- ☐ Yes – please call 715-221-6462 to report a new diagnosis
- ☐ No

MARSHFIELD CLINIC RESEARCH INSTITUTE
IN COLLABORATION WITH
SANFORD HEALTH
